# Multiomic insights into sucrose accumulation in sugarcane

**DOI:** 10.1101/2024.06.18.599623

**Authors:** Alexandre Hild Aono, Ricardo José Gonzaga Pimenta, Jéssica Faversani Diniz, Marishani Marin Carrasco, Guilherme Kenichi Hosaka, Fernando Henrique Correr, Ana Letycia Basso Garcia, Estela Araujo Costa, Alisson Esdras Coutinho, Luciana Rossini Pinto, Marcos Guimarães de Andrade Landell, Mauro Alexandre Xavier, Dilermando Perecin, Monalisa Sampaio Carneiro, Thiago Willian Balsalobre, Reginaldo Massanobu Kuroshu, Gabriel Rodrigues Margarido, Anete Pereira de Souza

## Abstract

Sugarcane (*Saccharum* spp.) holds significant economic importance in sugar and biofuel production. Despite extensive research, understanding highly quantitative traits, such as sucrose content, remains challenging due to the complex genomic landscape of the crop. In this study, we conducted a multiomic investigation to elucidate the genetic architecture and molecular mechanisms governing sucrose accumulation in sugarcane. Using a biparental cross (IACSP95-3018 × IACSP93-3046) and a genetically diverse collection of sugarcane genotypes, we evaluated the soluble solids (Brix) and sucrose content (POL) across various years and environments. Both populations were genotyped using a genotyping-by-sequencing (GBS) approach, with single nucleotide polymorphisms (SNPs) identified via bioinformatics pipelines. Genotype‒phenotype associations were established using a combination of traditional linear mixed-effect models and machine learning algorithms. Furthermore, we conducted an RNA sequencing (RNA-Seq) experiment on genotypes exhibiting distinct Brix and POL profiles across different developmental stages. Differentially expressed genes (DEGs) potentially associated with variations in sucrose accumulation were identified. All findings were integrated through a comprehensive gene coexpression network analysis. Strong correlations among the evaluated characteristics were observed, with estimates of modest to high heritabilities. By leveraging a broad set of SNPs identified for both populations, we identified several SNPs potentially linked to phenotypic variance. Our examination of genes close to these markers facilitated the association of such SNPs with DEGs in genotypes with contrasting sucrose levels. Through the integration of these results with a gene coexpression network, we delineated a set of genes potentially involved in the regulatory mechanisms of sucrose accumulation in sugarcane, collectively contributing to the definition of this critical phenotype. Our findings constitute a significant resource for biotechnology and plant breeding initiatives. Furthermore, our genotype‒phenotype association models hold promise for application in genomic selection, offering valuable insights into the molecular underpinnings governing sucrose accumulation in sugarcane.

## Introduction

Sugarcane holds significant importance in the global economy, particularly in terms of biofuel and sugar production (FAOSTAT 2023). Due to its remarkable capacity for sugar storage, sugarcane is the primary global source of sugar (Mirajkar et al. 2019). With the continual rise in sugar demand, there is a pressing need for the development of more productive varieties. Central to sugarcane breeding programs is the maximization of yield, measured in terms of sugar production per area (Cursi et al. 2022). This optimization encompasses resistance to abiotic and biotic stressors and several secondary traits, facilitating sugarcane cultivation across diverse environmental conditions.

Despite notable advancements in sugarcane varieties, the process of cultivar generation through breeding can last up to 12 years (De Morais et al. 2015). Sugarcane breeding typically involves three main stages: (i) creating genetic variability through controlled crosses; (ii) preliminary selection across numerous experiments with limited replicates; and (iii) advanced selection, with an adequate number of replicates and environments to enable precise selection (Gazaffi et al. 2015). Given the extensive time and costs associated with field evaluations, the integration of molecular-assisted technologies holds promise for accelerating breeding progress and increasing genetic gains, particularly regarding sucrose content, an aspect where sugarcane breeding progress remains slow (Chen et al. 2019). However, the intricate genomic complexity of sugarcane poses a challenge in understanding the genetic architecture underlying sugar accumulation and consequently hinders the development of effective molecular breeding efforts.

Modern sugarcane cultivars are derived from crosses between *Saccharum officinarum* (2n = 8x = 80, x = 10) (D’Hont et al. 1998) and *Saccharum spontaneum* (from 2n = 5x = 40 to 16x = 128, x = 8) (Panje and Babu 1960), followed by several backcrosses with *S. officinarum* to increase sucrose content (Cuadrado et al. 2004). While *S. spontaneum*, a wild sugarcane species, exhibits high stress resistance, it has a low sucrose content and abundant biomass (Mirajkar et al. 2019). Wild sugarcane can store approximately 2% of its fresh weight as sucrose, whereas the theoretical storage capacity of cultivated sugarcane can reach 27% (Bull and Glasziou 1963). Understanding the genetic mechanisms associated with these contrasting sugar accumulation profiles is challenging because of factors such as varying ploidy levels, frequent aneuploidies, and substantial cytogenetic complexity (Aono et al. 2021).

The quantitative trait loci (QTLs) associated with sugar-related traits exhibit a highly complex genetic architecture (Ming et al. 2002; Costa et al. 2016; Balsalobre et al. 2017), and there is limited information regarding the extent, effect, and genomic regions associated with phenotypic variability. This polygenic action encompasses diverse metabolic pathways and biological processes, particularly during the maturation phase, which dictates sucrose accumulation in mature sugarcane (Datir and Joshi 2016). Sucrose synthesis occurs primarily in sugarcane leaves. The sucrose is then transported through the phloem and stored in culms (Sachdeva et al. 2011). Its metabolism is regulated by diverse sucrose-synthesizing and hydrolyzing enzymes, including sucrose synthase, sucrose phosphate synthase, and invertases (Datir and Joshi 2016).

In addition to sucrose metabolism, other metabolic pathways, such as photosynthesis and carbon partitioning, influence sucrose accumulation rates in sugarcane (Sachdeva et al. 2011). Genes associated with stress responses also play significant roles in the efficiency of this mechanism, with notable implications for the regulatory actions of jasmonic acid, abscisic acid, ethylene, and gibberellin (Papini-Terzi et al. 2009). Therefore, integrative methodologies present considerable potential for dissecting these mechanisms and identifying critical regulatory elements involved in sucrose accumulation. Such endeavors are invaluable for biotechnology and molecular breeding approaches, especially given that the modification of genes associated with sucrose metabolism and transport has not yielded satisfactory outcomes (Qin et al. 2021).

Our study explored the intricate genetic architecture underlying sucrose accumulation in sugarcane. Through the integration of diverse omics datasets derived from a range of sugarcane genotypes, we not only offer insights into potential genotype-phenotype associations but also conduct a thorough exploration of how these associations influence the molecular mechanisms of sucrose accumulation. Leveraging a variety of methodologies, including linear mixed-effects modeling, machine learning algorithms, and gene coexpression networks, in addition to genomic analyses and differential expression gene comparisons, we elucidate crucial mechanisms and pivotal regulators governing this multifaceted process.

## Material and Methods

### Plant Material

Two distinct sugarcane populations were utilized in this study to investigate genotype‒phenotype associations. The first population (Pop1) comprised a panel of 97 diverse sugarcane accessions (Supplementary Table S1), and the second population (Pop2) consisted of 219 progeny genotypes derived from a cross between the elite clone IACSP953018 (female parent) and the commercial variety IACSP933046 (male parent). Both populations were developed by the Sugarcane Breeding Program at the Agronomic Institute of Campinas (IAC) in Ribeirão Preto, São Paulo, Brazil (4°52′34″ W, 21°12′50″ S). Planting occurred in 2013 for Pop1 and in 2011 for Pop2, following a complete block design with 4 blocks for Pop1 and 2 blocks for Pop2. In Pop1, three plants per experimental unit were planted in 1.5 m rows, with 0.5 m spacing between the plants. In Pop2, the plants were planted in 2 m rows with a spacing of 1.5 m between the plants. Additionally, the Pop1 experiment was replicated three times, corresponding to harvest times in May, July, and September (Coutinho et al. 2022).

Furthermore, three cultivars were selected for an RNA sequencing (RNA-Seq) experiment based on their divergent sugar content profiles. These genotypes were planted with three replicates in a field at the Federal University of São Carlos in Araras, São Paulo, Brazil (47°23′5″ W, 22°18′41″ S). Specifically, the selected genotypes included (i) IN84-58, a representative of *S. spontaneum* with low soluble solids content (Brix); (ii) the SP80-3280 hybrid, characterized by high Brix measurements; and (iii) the hybrid R570, which also exhibits high Brix measurements.

### Phenotyping

The genotypes from Pop1 and Pop2 were phenotyped for Brix and sucrose content (POL) following the methods described in Consecana (2006). For Pop1, evaluations were conducted in ratoon cane in 2014 and 2015, with one-year intervals between harvests. For Pop2, evaluations were conducted in plant cane in 2012 and in ratoon cane in 2013 and 2014.

Each trait in each population and replication was modeled using the following linear mixed-effects model:

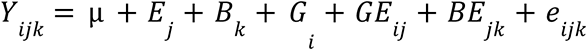

where *Y_ijk_* represents the phenotypic measurement of the *i*-th genotype in the *j*-th year and *k*-th block; µ is the overall trait mean; *E_j_* is the fixed effect of the *j*-th year; *B_k_* is the fixed effect of the *k*-th block; *G* is the random effect of the *i*-th genotype (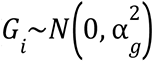, with 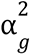 representing the genetic variance); *GE_ij_* is the random effect of the interaction between the *i*-th genotype and *j*-th year; *BE_jk_* is the random effect of the interaction between the *k*-th block and the *j*-th year; and *e_ijk_* is the residual term (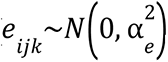, with 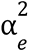 representing the residual variance). For Pop1, a separate model was created for each replication and combined with an additional model:

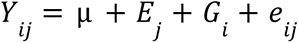

where *Y_ij_* represents the best linear unbiased predictors (BLUPs) for the genetic effects of the *i*-th genotype in the *j*-th harvest time; µ is the overall mean; *E_j_* is the fixed effect of the *j*-th harvest time; *G_i_* is the random effect of the *i*-th genotype; and *e_ij_* is the residual term. To estimate the variance components and BLUPs, we utilized the R package ASReml-R v4.1.0 (Butler et al. 2009) employing the restricted maximum likelihood (REML) approach. We assessed the significance of fixed effects using Wald tests and the significance of random effects using analyses of deviance and likelihood ratio tests (LRTs). Broad-sense heritabilities were estimated according to Cullis et al. (2006):

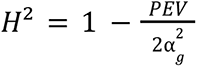

Where *PEV* represents the prediction error variance and 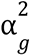 represents the estimated genetic variance. Additionally, we computed heritabilities based on the ratio of genotypic variance to total phenotypic variance.

To facilitate direct comparisons of estimates between populations, BLUP values were rescaled to the range of 0–1. Multivariate and descriptive analyses were performed using R statistical software v4.1.2 (R Core Team 2013). To assess phenotypic similarities between genotypes, we conducted complete linkage hierarchical clustering analysis based on Euclidean distances.

### Genotyping

The populations were genotyped using a genotyping-by-sequencing (GBS) approach following the methodologies outlined by Elshire et al. (2011) and Poland et al. (2012). A total of 94 individuals from Pop1 and 182 individuals from Pop2, consisting of 180 progeny genotypes and their respective parents, were genotyped. In Pop1, genotyping was accomplished utilizing a combination of the restriction enzymes *PstI* and *MseI*, as described by Pimenta et al. (2021). In Pop2, only the enzyme *PstI* was employed, following the methodology described by Aono et al. (2020). Sequencing procedures were performed using the Illumina platform, with the NextSeq 500 platform utilized for Pop1 and a combination of the GAIIx and NextSeq 500 platforms for Pop2.

Single nucleotide polymorphism (SNP) calling was executed utilizing the TASSEL-GBS pipeline (Glaubitz et al. 2014), which was adapted for polyploid species (TASSEL4-POLY, Pereira et al. 2018). Read mapping was conducted utilizing the sugarcane genome sequence of the cultivar SP70-1143 obtained through methylation filtration (Grativol et al. 2014) and the Bowtie2 v2.2.5 tool (Langmead and Salzberg 2012). The selection of the sugarcane genomic reference was based on its demonstrated superiority in handling sugarcane GBS data, as evidenced by Aono et al. (2020). For subsequent analyses, biallelic SNPs were selected based on the following stringent criteria: (i) a minimum depth of 50 reads per individual at a SNP position, (ii) a minimum allele frequency of 10%, and (iii) a maximum of 10% missing data. Due to the aneuploid nature of the sugarcane genome, SNPs were organized based on allele proportions, representing the ratio between the number of reads of the reference allele and the total number of reads. The genotypic data were subjected to multivariate analysis using uniform manifold approximation and projection (UMAP) for dimension reduction, implemented with the R package Umap v0.2.10.0 (McInnes et al. 2018).

### RNA Sequencing and Transcriptome Analyses

The culm samples were collected from the +1 internode of the selected genotypes at development times of 6, 8, 10, and 12 months. We employed three biological replicates and three technical replicates. RNA-Seq libraries were prepared and sequencing was performed on the HiSeq 2500 Illumina platform following the protocol described by Hosaka et al. (2021).

Raw sequencing reads were filtered using Trimmomatic v0.39 (Bolger et al. 2014). This process involved removing base pairs with quality scores below 3 at the beginning and end of the reads, excluding regions with an average quality less than 20 in a window of 4 base pairs, and discarding reads shorter than 75 base pairs. The filtered reads were then aligned to the genomic references of *S. officinarum* (GenBank GCA_020631735.1) and *S. spontaneum* (GenBank GCA_022457205.1) using STAR v2.7.3 (Dobin and Gingeras 2015). Each allele was considered an independent reference, resulting in 12 alignments per sequencing file—eight for *S. officinarum* and four for *S. spontaneum*. The aligned reads were sorted based on genomic positions using SAMtools v1.12 (Li et al. 2009), and transcriptomes were assembled using Stringtie v2.1.6 (Pertea et al. 2015).

To reduce redundancy in the assembled transcriptomes, we utilized CD-HIT (Fu et al. 2012). First, redundancies within each species were eliminated by combining individual allele transcriptomes. Then, the combined transcriptomes of both species were merged to generate the final transcriptome, which was also evaluated with CD-HIT (Fu et al. 2012) for redundancy removal. This approach ensured the selection of a single representative transcript for each set of sequences, facilitating downstream analyses.

Transcriptome assembly was evaluated using the BUSCO v5.5.0 tool (Simão et al. 2015), and the results were compared against those of both the Viridiplantae and Eukaryota databases. Transcript annotation was performed utilizing Trinnotate v4.0.1 software (Griffith et al. 2015) and the UniProt database (UniProt Consortium 2019).

To obtain gene expression estimates, we utilized Salmon v1.9.0 software (Patro et al. 2017). Following the quantification of gene expression across samples, we implemented additional filtering steps to establish a refined set of gene expression estimates for subsequent analyses. Specifically, we computed the gene counts per million (CPM) values using the edgeR v3.36.0 package (Robinson et al. 2010), ensuring that only genes with at least three samples possessing a minimum of 10 CPMs were retained.

The distribution of RNA-Seq samples was visualized through a scatter plot generated from a principal component analysis (PCA) conducted on the gene expression estimates using R statistical software v4.1.2 (R Core Team 2013).

### Genotype-Phenotype Associations

To identify associations between SNPs and the phenotypic values of Brix and POL, we employed two different approaches: a genome-wide association study (GWAS) and machine learning techniques (Aono et al. 2022). For the GWAS, we used the R package ASReml-R v4.1.0 (Butler et al. 2009) with the REML approach. We modeled the BLUPs estimated from the previous models (*Y_i_*) of the *i*-th individual using a linear mixed-effects model for each *k*-th SNP:

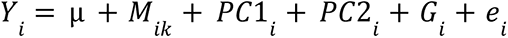

where µ is the overall mean; *M_ik_* is the fixed effect associated with the allele proportion of the *k*-th SNP of the *i*-th individual; *PC*1*_i_* and *PC*2*_i_* are fixed effects associated with the first components of the *i*-th individual estimated through a principal component analysis (PCA) performed with the SNP data (missing values were imputed as the mean of the observed values for each SNP); *G_i_* is the random polygenic effect of the *i*-th individual (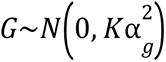, with *K* representing the genomic relationship matrix and 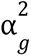 representing the genetic variance); and *e_i_* is the residual term (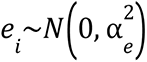, with 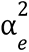 representing the residual variance). We also employed the same model to investigate potential associations between SNP markers and phenotypic measures within the biparental population (Pop2). However, we opted to exclude the contributions of *PC*1_*i*_ and *PC*2_*i*_ due to the inherent similarities observed among individuals.

We calculated the genomic relationship matrix *K* using the R package AGHmatrix v2.1.0 (Amadeu et al. 2016) with a fixed ploidy of 10 and allele dosages calculated based on ten intervals with a discretization length of 0.1 in the allele proportions. We evaluated the significance of each SNP using a Wald test with Bonferroni and false discovery rate (FDR) corrections, setting an adjusted Bonferroni p value threshold of 0.05 for considering a SNP associated with a phenotype.

As the genomic reference utilized for SNP calling is assembled at the scaffold level (Grativol et al. 2014), we employed an additional step to incorporate markers in linkage disequilibrium (LD) with the SNPs identified through GWAS. For each SNP associated with the traits under investigation, Pearson correlations were conducted with the remaining set of markers. We identified potential associations by selecting markers with a minimum absolute Pearson correlation coefficient of 0.75 and a significance threshold of p ≤ 0.05. The representation of these associations was constructed utilizing a correlation graph generated with the R package igraph v1.3.5 (Csardi and Nepusz 2006).

For the machine learning approach, we used feature selection (FS) techniques implemented in Python v3.10.12 with the scikit-learn v1.2.2 library (Pedregosa et al. 2011). Each SNP represented a feature, and the BLUP value was the target to be predicted. We employed three algorithms: gradient tree boosting (GTB), L1-based FS with a linear support vector regression system (SVM), and Pearson correlation (with a p value cutoff of 0.05). An SNP was considered to have a phenotypic association if it was identified by all three methods simultaneously (Aono et al. 2022).

In addition, to evaluate genotype‒phenotype associations identified via FS techniques, we compared the predictive performance of genomic prediction models trained using the complete set of SNPs against models trained exclusively with FS-selected markers, employing a leave–one-out cross-validation methodology. We employed two machine learning algorithms implemented in Python v3.10.12 with the scikit-learn v1.2.2 library (Pedregosa et al. 2011) as our modeling approach: support vector regression (SVM) and adaptive boosting (AdaBoost). Model accuracies were evaluated using Pearson correlation coefficients and mean squared errors, measured considering the observed and predicted phenotypic values.

To elucidate the potential functional implications of the identified mutations, we associated all SNPs identified in correlation with Brix or POL measures with potential gene sequences retrieved from the assembled transcriptome. Specifically, we conducted an alignment of all assembled transcripts with the sugarcane genome sequence of the cultivar SP70-1143 using the BLASTn v2.11.0+ tool (Altschul et al. 1990). For each SNP-associated scaffold, we considered a maximum of 5 alignments, applying an E-value cutoff of 1e-6.

Based on the alignments obtained, we performed gene ontology (GO) enrichment analyses using the R package topGO v2.46.0 (Alexa and Rahnenführer 2009). We established an FDR-adjusted p value threshold of 0.05 to determine the significance of GO term enrichment. All enriched GO categories were summarized using the Revigo tool (Supek et al. 2011).

### Differential Gene Expression and Coexpression Networks

The identification of differentially expressed genes (DEGs) was conducted using the filtered gene set and the R package DESeq2 v1.34.0 (Love et al. 2014). To identify genes potentially associated with differences in sugar accumulation profiles, we compared the gene expression profiles of the IN84-58 genotype (low sugar accumulation) with those of the hybrids SP80-3280 and R570 (high sugar accumulation) considering the developmental time point as a factor in a model fitted according to a factorial design. Additionally, these sets of DEGs were intersected with contrasts performed on developmental time points of SP80-3280 and R570 gene expression estimates. An FDR-adjusted p value threshold of 0.05 and a log2-fold change of 1.5 were applied to define DEGs.

GO enrichment analysis was conducted using the R package topGO v2.46.0 (Alexa and Rahnenführer 2009), with an FDR-adjusted p value cutoff of 0.05. All enriched GO categories were summarized using the Revigo tool (Supek et al. 2011).

Using gene expression estimates organized in transcripts per million (TPM), we constructed a gene coexpression network employing the weighted gene coexpression network analysis (WGCNA) method implemented in the R package WGCNA v1.72.1 (Langfelder and Horvath 2008). Initially, we determined the soft power parameter (β) by selecting the value that resulted in a minimum R² of 0.8 and maximum mean connectivity, ensuring that the network approximated a scale-free topology. Subsequently, based on Pearson correlation coefficients and the estimated β, we computed an adjacency matrix, which was then used to define a dissimilarity matrix derived from a calculated topological overlap matrix. Finally, average-linkage hierarchical clustering was applied to the dissimilarity matrix, and adaptive branch pruning was performed to identify modules of coexpressed genes.

GO module enrichment analysis was performed using the R package topGO v2.46.0 (Alexa and Rahnenführer 2009) with an FDR-adjusted p value cutoff of 0.05.

### Multiomics Analyses

To integrate the findings from various analyses, we conducted a comprehensive investigation using a gene coexpression network model. Initially, we examined each network module based on the following criteria: (i) the number of genes associated with GWAS/LD results, (ii) the number of genes associated with FS results, and (iii) the number of DEGs identified in intersection contrasts.

Based on these criteria, we selected groups of coexpressed genes and constructed specific gene coexpression networks for the IN84-58, SP80-3280, and R570 genotypes using the highest reciprocal rank (HRR) approach (Mutwil et al. 2010). We utilized gene expression estimates organized in TPMs for genes within these groups and generated a Pearson correlation coefficient matrix. Subsequently, we constructed the network by considering the 30 strongest absolute correlations (minimum R Pearson correlation of 0.7) and modeling a graph using the R package igraph v1.3.5 (Csardi and Nepusz 2006). Furthermore, we evaluated the network architecture using different centrality measures for each gene, including degree, Kleinberg’s hub score, and betweenness.

## Results

### Phenotyping and Genotyping

Brix and POL were analyzed through linear mixed effects models to comprehensively assess variance components and estimate the genetic contributions of the evaluated phenotypes (Supplementary Table S2). Notably, substantial correlations were detected between these traits in both populations studied, with Pearson correlation coefficients of 0.95 for the 97 sugarcane accessions (Pop1) and 0.9 for the 219 progeny genotypes resulting from the biparental cross (Pop2). Upon employing BLUP estimates (Supplementary Table S3), the correlation coefficient in Pop1 decreased to approximately 0.9, but in Pop2, it increased to approximately 0.93 (Fig. 1a and b). This divergence in correlations highlights potential environmental influences that may have been captured by the preceding correlation analyses.

**Fig. 1.**
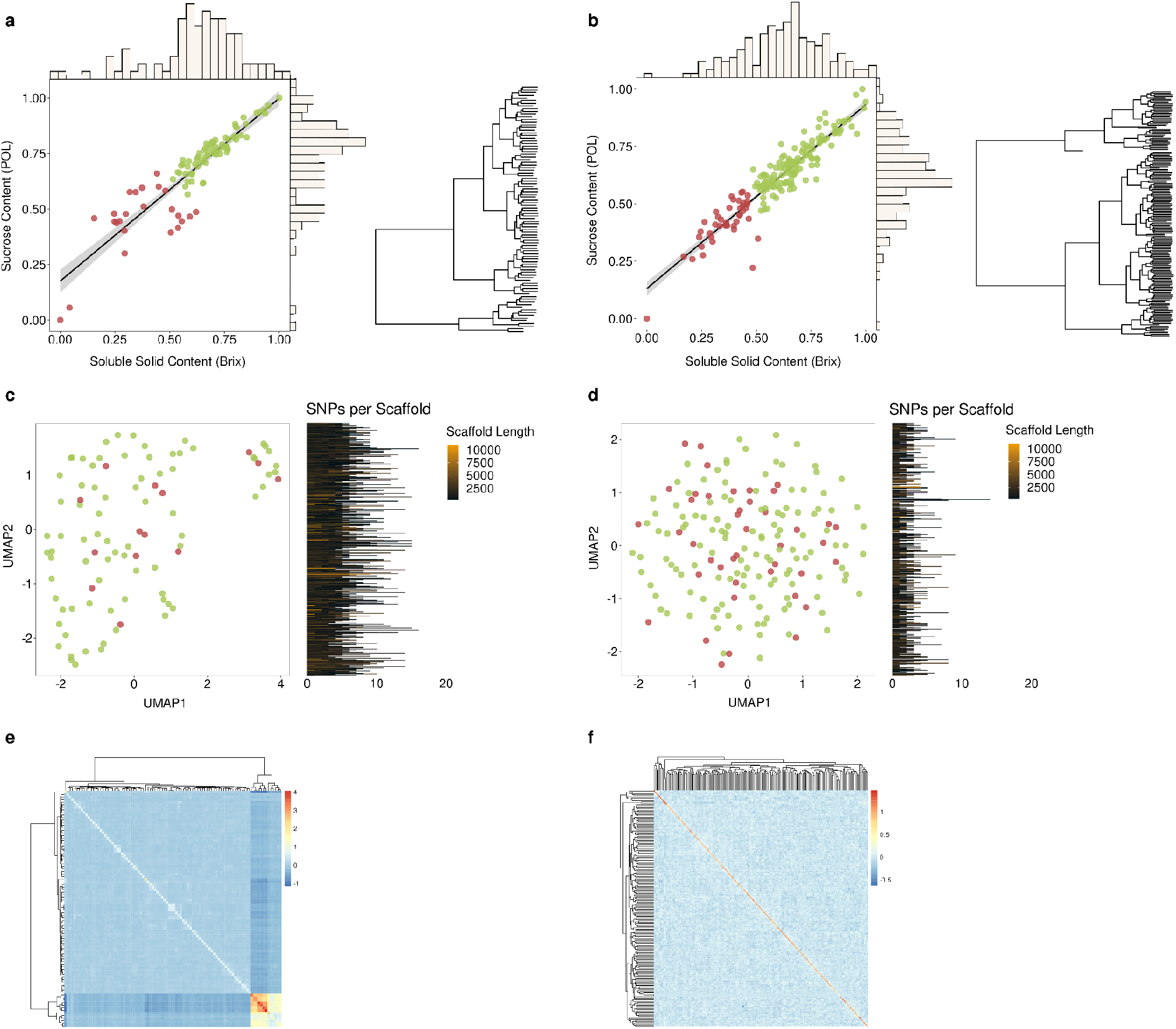
Distribution of genotypic data and best linear unbiased predictions (BLUPs) for soluble solids content (Brix) and sucrose content (POL) in two evaluated populations: Pop1, consisting of a panel of 97 sugarcane accessions; and Pop2, comprising 219 progeny genotypes derived from a biparental cross. Scatter plots illustrating associations between Brix and POL are depicted for Pop1 (a) and Pop2 (b), along with dendrograms illustrating clustering profiles for each population. Additionally, uniform manifold approximation and projection (UMAP) analyses are presented for Pop1 (c) and Pop2 (d) based on SNP data. Individuals are colored according to a hierarchical clustering analysis of the phenotypic measures. Genomic relationship matrices are provided for Pop1 (e) and Pop2 (f), indicating the genetic relationships within each population.

Estimates of broad-sense heritability using variance ratios were greater in Pop1, with values ranging from approximately 0.89 for Brix in experimental unit 2 to approximately 0.97 for POL in experimental unit 3. Heritability estimates obtained through Cullis’ method (Cullis et al. 2006) were consistent with the values observed for the ratios, differing by approximately 1%. In contrast, Pop2 exhibited lower estimates (∼0.36 for Brix and ∼0.37 for POL). The higher estimates in Pop1 can be attributed to the more pronounced phenotypic variation among individuals in the panel, as Pop1 includes commercial sugarcane cultivars from Brazilian breeding programs as well as *S. spontaneum* and *S. robustum* accessions, representing traditional energy cane clones. Remarkably, the highest estimates of genetic effects for Brix and POL were observed for IACCTC059552, a modern sugarcane hybrid, and the lowest were recorded for IACBIO275, an energy cane clone (Supplementary Table S3).

The genetic differences observed in the populations and models were found to be statistically significant (Supplementary Table S2). In the biparental population (Pop2), clear evidence of heterosis was observed, with a significant proportion of progeny genotypes exhibiting estimates larger than those of the most productive parent (21 individuals for Brix and 26 for POL). There were no significant interactions detected between genetic and year effects.

Hierarchical clustering analysis of the phenotypic measures from both populations revealed a distinct separation of genotypes into two groups, colored in green and red in Fig. 1a and b. By contrasting the Brix and POL measures between these groups, statistically significant differences were identified through t tests. The p values for Brix in Pop1 and Pop2 were 1.19e-13 and < 2.2e-16, respectively, and for POL in Pop1 and Pop2, the p values were 3.014e-07 and < 2.2e-16, respectively.

The sequencing of the GBS libraries generated a substantial amount of data, with 863,889,004 reads for Pop1 and 1,103,163,250 reads for Pop2. Subsequent analysis using the TASSEL-POLY pipeline identified 874,597 and 137,757 SNPs for Pop1 and Pop2, respectively. To ensure data reliability, rigorous filtering criteria were applied, resulting in a final set of 16,166 SNPs for Pop1 and 2,178 SNPs for Pop2 (Supplementary Tables S4 and S5).

Multivariate analysis (Fig. 1c and d) did not reveal any distinct patterns correlating genotypes with phenotypes, suggesting challenges in elucidating the genetic architecture underlying the observed traits. In Pop2, the absence of genotypic clusters was consistent with expectations due to the crossing nature of the genotypes. Conversely, in Pop1, a discernible pattern emerged, possibly indicating a subgroup of individuals with closer genetic relatedness, although this pattern did not correspond to any observed associations with sugar-related phenotypes. Similar patterns were also observed in the genomic relationship matrices (Fig. 1e and f), further supporting the existence of a distinct subgroup within Pop1.

### Transcriptome Assembly and Gene Expression Estimates

The RNA-Seq experiment generated a substantial dataset consisting of 1,240,508,982 paired-end sequencing reads, each with a length of 100 base pairs. The mean number of reads per sample was 11,486,194.28 (Supplementary Table S6). Following stringent filtering procedures, 1,046,816,212 paired-end sequencing reads were retained, accounting for approximately 84.39% of the initial reads.

Subsequently, the filtered reads were aligned to the genomes of *S. spontaneum* and *S. officinarum* and assembled at the allele level, facilitating independent assemblies for each species allele. The transcript quantities assembled for each allele of *S. spontaneum* were as follows: A) 53,826, B) 53,524, C) 52,249, and D) 52,569. For *S. officinarum*, the quantities were A) 55,272, B) 53,563, C) 53,809, D) 50,945, E) 49,668, F) 46,037, G) 44,220, and H) 39,048.

To minimize redundancy and streamline the dataset, the transcripts assembled per allele in each species were combined, and CD-HIT software was utilized. This process resulted in the generation of 138,774 transcripts for *S. spontaneum* and 201,646 transcripts for *S. officinarum*. Subsequently, by combining these two transcriptomes and applying CD-HIT, a final comprehensive transcriptome comprising 291,959 transcripts was obtained. This integrated approach not only established a comprehensive transcriptome reference for both species but also facilitated the determination of the origin of each gene, enabling further evolutionary inferences to be made.

The transcriptome assembly strategy generated transcripts with sizes ranging from 99 to 16,513 base pairs, with 291,615 transcripts (∼99.88%) presenting sizes greater than 200 nucleotides (the transcript N50 length was 1,765 bp). A comparison of these transcripts with the Eukaryota and Viridiplantae databases using BUSCO software revealed that 99.6% (86.3% of duplicated associations) and 99.7% (83.5% of duplicated associations) of the sequences were complete, respectively. Due to the use of allele-specific genome references for assembly, we expected a high percentage of duplications to be observed.

We identified a set of 46,098 genes by selecting those with at least three samples presenting 10 CPMs, and these genes were subsequently used for further analyses. Gene annotations were obtained through comparisons with the UniProt database, resulting in successful alignment of all genes with UniProt proteins. This facilitated the retrieval of diverse annotations for functional analyses. Specifically, 37,196 genes (∼80.69%) were found to correspond to GO terms. Analysis of the gene expression data using PCA revealed a distinct dispersion pattern across samples, effectively separating the genotypes (Fig. 2a). Notably, the IN84-58 genotype, representing *S. spontaneum*, exhibited more pronounced differences than the other genotypes.

**Fig. 2.**
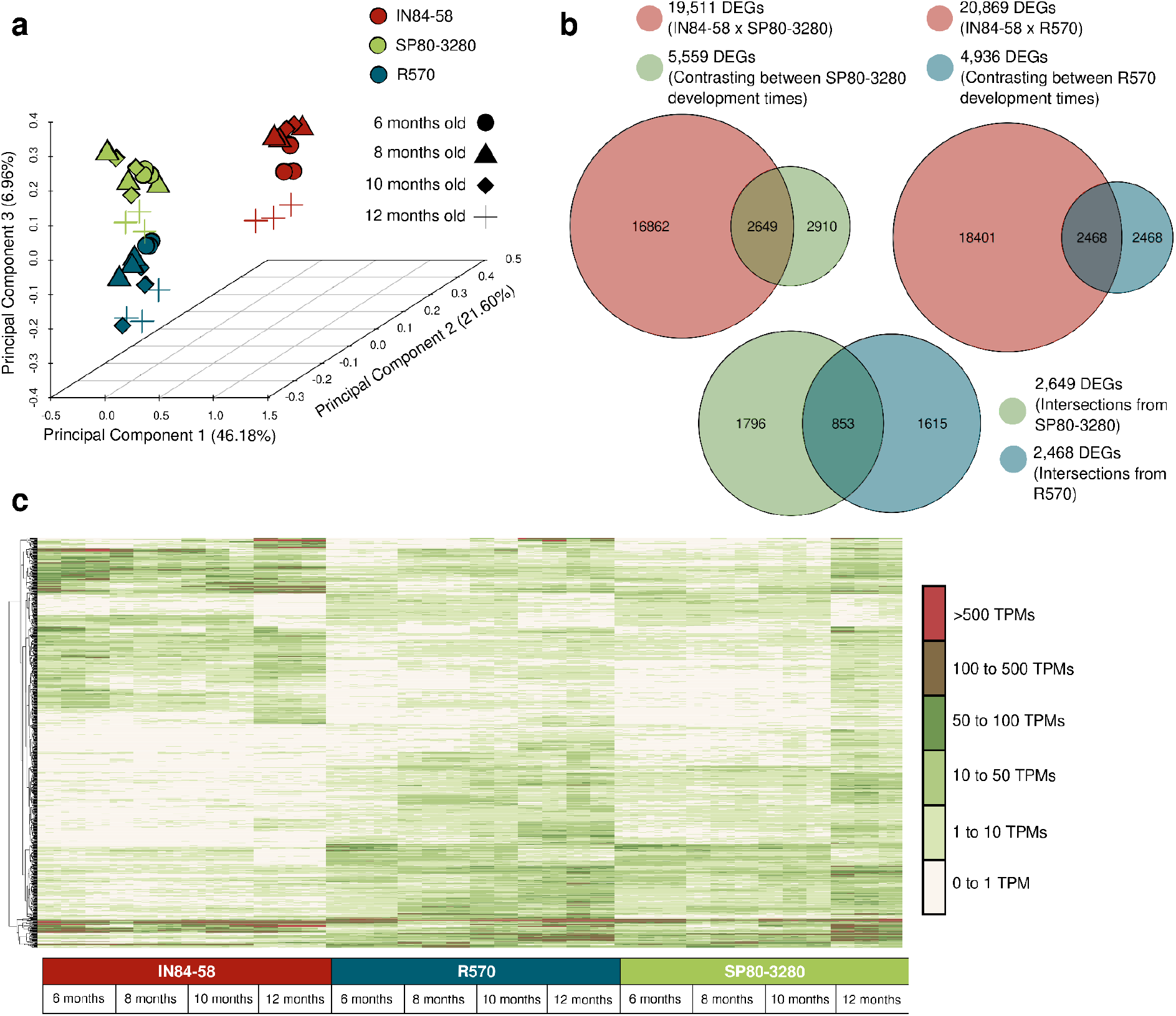
Gene expression analyses. (a) Principal component analysis (PCA) showing gene expression patterns across developmental time points (6, 8, 10, and 12 months old) for the IN84-58, SP80-3280, and R570 genotypes. (b) Identification of differentially expressed genes (DEGs) through intergenotype comparisons. (c) Heatmap illustrating the expression profiles of the final set of 853 DEGs selected for analysis.

### Genotype-Phenotype Associations

In our study aimed at identifying genotype‒phenotype associations, we initially employed a linear mixed-effects model to conduct the GWAS analysis (Table 1). Consistent with our expectations, the analysis revealed a greater number of associations in Pop1 than in Pop2, which was attributed to the greater genetic variability observed within Pop1. Specifically, in Pop1, we identified 7 SNPs significantly associated with Brix measures and 6 SNPs significantly associated with POL. Notably, 5 SNPs exhibited simultaneous associations with both phenotypes, which aligns with the anticipated outcome due to the pronounced correlation between Brix and POL (Fig. 1a). Conversely, fewer associations were observed in Pop2, with only 1 SNP associated with Brix and another 1 associated with POL. Subsequent examination of the allelic proportion profiles of these SNPs in comparison to the phenotypic measurements revealed a consistent distribution pattern (Fig. 3a and b), supporting the validity of the observed associations.

**Fig. 3.**
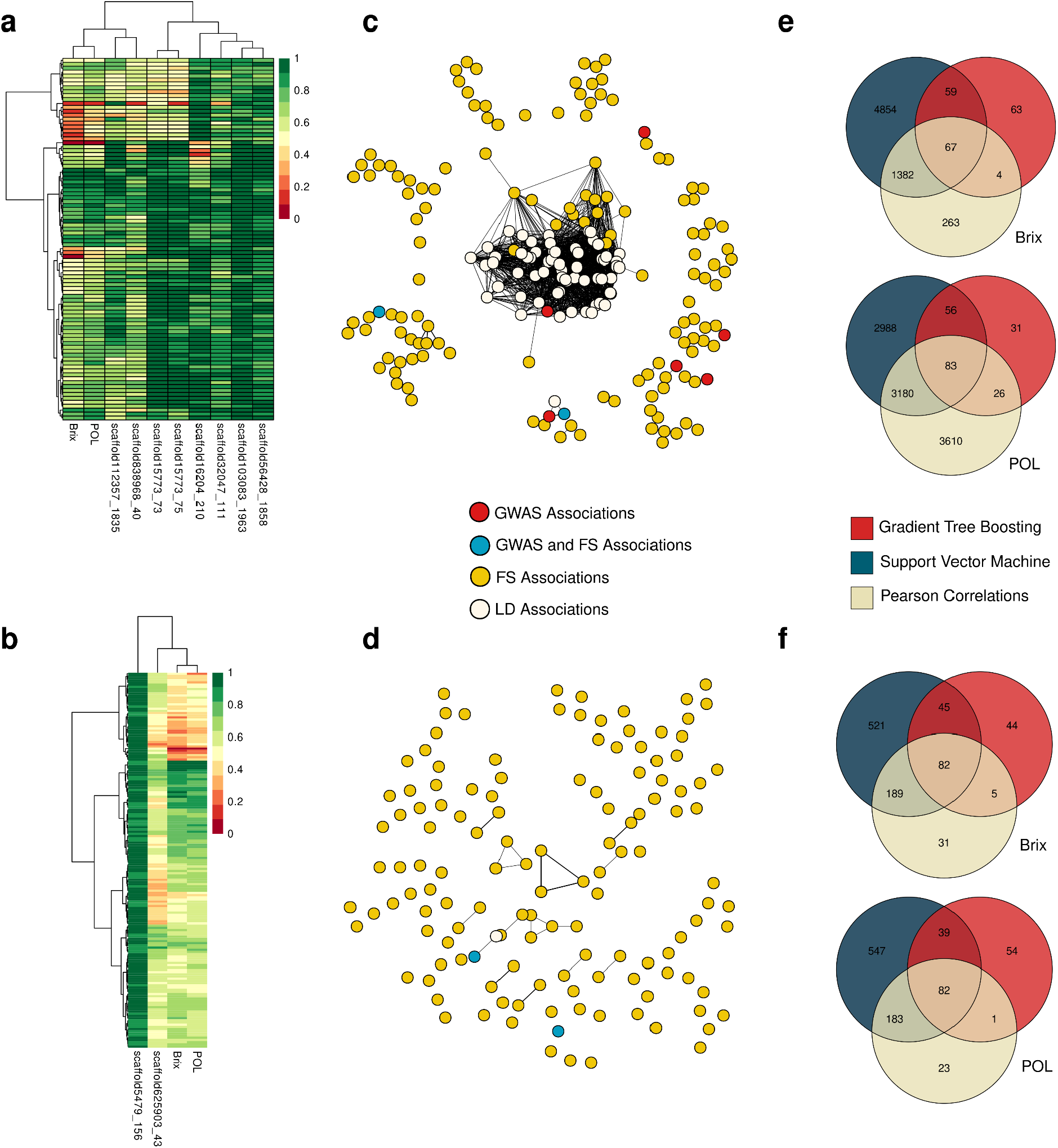
Allelic proportions of single nucleotide polymorphisms (SNPs) identified through a genome-wide association study (GWAS) related to soluble solids content (Brix) and sucrose content (POL) in two populations: Pop1, comprising a panel of 97 sugarcane accessions (a); and Pop2, consisting of 219 progeny genotypes derived from a cross between the elite clone IACSP953018 (female parent) and the commercial variety IACSP933046 (male parent) (b). Linkage disequilibrium (LD) networks for Pop1 (c) and Pop2 (d) constructed based on the associations of SNPs identified through GWAS with the remaining markers in the dataset. SNPs were selected using feature selection (FS) techniques, including gradient tree boosting (GTB), L1-based FS employing linear support vector regression (SVM), and Pearson correlations (with a p value threshold of 0.05), in Pop1 (e) and Pop2 (f).

**Table 1.** Genome-wide association study (GWAS) results for soluble solids content (Brix) and sucrose content (POL) across two distinct populations: Pop1, comprising a panel of 97 sugarcane accessions; and Pop2, consisting of 219 progeny genotypes derived from a cross between the elite clone IACSP953018 (female parent) and the commercial variety IACSP933046 (male parent). Adjusted p values were calculated using both Bonferroni and false discovery rate (FDR) corrections. SNPs with Bonferroni-adjusted p values ≤ 0.05 were deemed to be significantly associated.

| Population | Trait | SNP | P value | FDR | Bonferroni |
| --- | --- | --- | --- | --- | --- |
| Pop1 | Brix | scaffold16204\|size3850_210_A/T | 2.88E-09 | 4.66E-05 | 4.66E-05 |
|  |  | scaffold15773\|size3862_73_C/A | 8.18E-08 | 0.0006490370472 | 0.001322046039 |
|  |  | scaffold838968\|size239_40_G/A | 1.20E-07 | 0.0006490370472 | 0.001947111141 |
|  |  | scaffold15773\|size3862_75_C/T | 1.67E-07 | 0.000673431525 | 0.0026937261 |
|  |  | scaffold112357\|size2063_1835_C/A | 5.35E-07 | 0.00172989985 | 0.00864949925 |
|  |  | scaffold32047\|size2236_111_G/T | 1.80E-06 | 0.004842866611 | 0.02905719966 |
|  |  | scaffold103083\|size2070_1963_G/T | 2.83E-06 | 0.006542895698 | 0.04580026989 |
|  | POL | scaffold16204\|size3850_210_A/T | 1.16E-09 | 1.88E-05 | 1.88E-05 |
|  |  | scaffold838968\|size239_40_G/A | 2.32E-08 | 0.000187183688 | 0.0003743673761 |
|  |  | scaffold15773\|size3862_73_C/A | 1.18E-07 | 0.0005698556482 | 0.00190506238 |
|  |  | scaffold15773\|size3862_75_C/T | 1.41E-07 | 0.0005698556482 | 0.002279422593 |
|  |  | scaffold32047\|size2236_111_G/T | 2.19E-06 | 0.007079909011 | 0.03539954505 |
|  |  | scaffold56428\|size2092_1858_A/G | 2.81E-06 | 0.007208389401 | 0.04544528141 |
| Pop2 | Brix | scaffold625903 \| size288_43_C/A | 1.42E-05 | 0.008841259904 | 0.03098643235 |
|  | POL | scaffold5479 \| size4842_156_C/T | 2.10E-05 | 0.03007418539 | 0.04568349979 |

Among the 10 SNPs identified, we retrieved annotations for only 4 SNPs (Supplementary Table S7). Among these SNPs, 2 were simultaneously associated with the Brix and POL traits in Pop1: an SNP at position 210 on scaffold16204 and an SNP at position 111 on scaffold32047. These SNPs corresponded to 5 genes annotated for anion transporters (gene_32017, gene_34208, gene_34382, gene_38431, and gene_39421) and 5 genes annotated for the protein FAR1 (gene_11104, gene_4861, gene_5373, gene_6529, and gene_8883). Another SNP associated with POL in Pop1 was located at position 1858 on scaffold56428 and annotated for 2 genes encoding serine/threonine-protein kinases (gene_34982 and gene_43850). The final annotated SNP was found in Pop2. It was located at position 156 on scaffold5479 and was associated with 2 genes encoding pentatricopeptide repeat-containing proteins (gene_32532 and gene_51857).

Of these 14 genes identified, 8 were exclusively found in *S. officinarum* (gene_32017, gene_34208, gene_38431, gene_39421, gene_11104, gene_4861, gene_5373, and gene_32532), 3 were found in both species (gene_34382, gene_6529, and gene_43850), and 3 were exclusively found in *S. spontaneum* (gene_8883, gene_51857, and gene_34982). Notably, most of the genes found in regions associated with contrasting sugar accumulation profiles are from the *S. officinarum* genome.

Regarding the GO terms associated with these GWAS-identified markers, we recovered a total of 27 GO terms (Supplementary Table S8). The most prominent GO terms were “regulation of transcription, DNA-templated” in the biological process category, “nucleus” in the cellular component category, and “zinc ion binding” in the molecular function category, and all of these terms were associated with 9 genes. These results indicate the potential role of these genes in the genetic regulation associated with differences in Brix and POL measurements.

Given that the genomic reference used lacked chromosome-level assembly, we implemented an alternative strategy to identify LD associations with the markers identified through GWAS. Utilizing pairwise Pearson correlations among allelic proportions, we identified 71 additional markers (Fig. 3c and d; Supplementary Table S9). Notably, only one marker was detected for Pop2, and this marker was specifically associated with the POL phenotypic trait. Conversely, the remaining 70 markers were correlated with GWAS-defined SNPs within Pop1. Of particular interest, 68 out of the 70 associations in Pop1 were associated with a single SNP (position 210 on scaffold16204), which was organized into smaller clusters across different scaffolds. For instance, SNPs located at positions 1216, 1265, 1268, 1270, 1271, and 1272 on scaffold 24635 exhibited correlations of approximately 0.8 with GWAS-defined SNPs. Similarly, SNPs located at positions 157, 166, 169, 173, and 199 on scaffold 562126 displayed correlations of approximately −0.8 with GWAS-defined SNPs. Such patterns suggest the presence of a coherent cluster of markers within the same QTL region, which may not have been adequately captured due to limitations in the genomic reference utilized.

We identified 75 additional genes associated with the LD markers (Supplementary Table S7). Interestingly, we observed no overlap between the genes identified through GWAS and LD analysis. However, we found annotations related to members of the kinase family in both sets of genes. Additionally, our analysis revealed novel annotations for various genes, including those encoding the E3 ubiquitin-protein ligase, the photosynthetic NDH subunit of subcomplex B3, the cleavage stimulation factor, and several transcription factors, such as MYB36, MYB87, RAX1, RAX2, and RAX3.

Through an evaluation of the GO terms associated with the genes surrounding the LD-associated markers, we identified a total of 197 terms (Supplementary Table S8). Prominent among the cellular components was the nucleus, which was associated with 28 genes. The most conspicuous molecular function was ATP binding, which was linked to 17 genes, and the prominent biological process was embryo sac development, which was correlated with 15 genes. Furthermore, several other noteworthy terms emerged, such as gene silencing by RNA, the cellular response to glucose stimulus, the regulation of glucose-mediated signaling pathway, the regulation of gene expression, carbohydrate transport, and the cellulose catabolic process.

By conducting an enrichment analysis combining GO terms associated with the GWAS and LD results, we identified 16 enriched biological process terms and 8 enriched molecular function terms (Supplementary Table S10). Our analysis highlighted regulatory processes such as the regulation of glucose-mediated signaling pathways, embryonic development, the negative regulation of DNA-templated transcription, and the positive regulation of abscisic acid-activated signaling pathways.

Moreover, by employing the established FS techniques, we successfully identified potential genotype‒phenotype associations (Supplementary Table S11; Fig. 3e and f). By applying a consensus approach involving the selection of markers identified by all three evaluated algorithms, we identified a total of 67 and 83 markers associated with the Brix and POL traits, respectively, in Pop1, with 15 overlapping SNPs. In Pop2, we identified a total of 82 markers associated with both the Brix and POL phenotypes, with an intersection of 15 SNPs. While no overlapping SNPs were observed between the populations, there were evident intersections among the FS methods for both phenotypic traits (Table 2).

**Table 2.** Single nucleotide polymorphisms (SNPs) associated with soluble solids content (Brix) and sucrose content (POL) were identified through the following feature selection strategies: gradient tree boosting (GTB), L1-based FS employing linear support vector regression (SVM), and Pearson correlation (with a p value threshold of 0.05). The populations employed were Pop1, consisting of a panel of 97 sugarcane accessions, and Pop2, consisting of 219 progeny genotypes derived from a cross between the elite clone IACSP953018 (female parent) and the commercial variety IACSP933046 (male parent).

| Population | Trait | Brix | POL | Intersection (Brix and POL) |
| --- | --- | --- | --- | --- |
| Pop1 | GTB | 193 | 193 | 25 |
|  | SVR | 6,362 | 6,307 | 5,632 |
|  | Pearson | 1,716 | 6,899 | 1,280 |
|  | Intersection (GTB, SVR, and Pearson) | 67 | 83 | 15 |
| Pop2 | GTB | 176 | 176 | 50 |
|  | SVR | 837 | 851 | 662 |
|  | Pearson | 307 | 289 | 205 |
|  | Intersection (GTB, SVR, and Pearson) | 82 | 82 | 31 |

To evaluate the impact of FS-selected SNPs on the phenotypic variation of Brix and POL, we employed a genomic prediction approach. Specifically, we assessed the predictive accuracies of these SNPs for Brix and POL and compared them with those obtained using the entire marker set. Employing a leave-one-out cross-validation methodology, we observed significant improvements in prediction accuracies with the FS-selected SNP set (Table 3). When utilizing the complete SNP set, the Pearson correlation coefficients between the observed and predicted values ranged from approximately 0.326 to 0.592 in Pop1 and from approximately 0.249 to 0.763 in Pop2. These accuracies were substantially enhanced upon subsetting the SNP dataset, yielding values ranging from approximately 0.760 to 0.811 in Pop1 and from approximately 0.661 to 0.846 in Pop2.

**Table 3.** Performance evaluation of machine learning algorithms (support vector regression (SVR) and adaptive boosting (AdaBoost)) for predicting soluble solids content (Brix) and sucrose content (POL) using genotype data. The performances of utilizing the entire SNP dataset (All) and employing feature selection (FS) are compared.

| Pop | Algorithm | Brix |  |  |  | POL |  |  |  |
| --- | --- | --- | --- | --- | --- | --- | --- | --- | --- |
|  |  | Pearson Correlation Coefficient |  | Mean Squared Error |  | Pearson Correlation Coefficient |  | Mean Squared Error |  |
|  |  | All | FS | All | FS | All | FS | All | FS |
| Pop1 | SVR | 0.546 | 0.806 | 0.036 | 0.017 | 0.592 | 0.760 | 0.023 | 0.015 |
|  | AdaBoost | 0.326 | 0.786 | 0.042 | 0.024 | 0.547 | 0.811 | 0.023 | 0.013 |
| Pop2 | SVR | 0.763 | 0.835 | 0.020 | 0.010 | 0.748 | 0.846 | 0.016 | 0.008 |
|  | AdaBoost | 0.253 | 0.668 | 0.031 | 0.021 | 0.249 | 0.661 | 0.023 | 0.016 |

We observed overlaps between findings from FS and GWAS coupled with LD analysis. For Brix in Pop1, we identified two SNPs by both approaches: one located in scaffold15773 at position 73, and one in scaffold112357 at position 1835. In Pop2, the markers identified by GWAS were also identified through FS. Remarkably, we further detected two additional markers situated within the same scaffolds identified by GWAS but not highlighted by LD tests. We speculate that these associations went unnoticed previously due to the rigorous parameters applied in our investigation. These SNPs were associated with both Brix and POL traits in Pop1, with one located in scaffold838968 at position 30 (identified at position 40 by GWAS) and one in scaffold15773 at position 3490 (reported at positions 73 and 75 by GWAS). These findings underscore the complementary nature of the methodologies employed in our study, reinforcing the validity of our results.

From the 238 SNPs identified using FS, we recovered 441 genes (Supplementary Table S7). Notably, when comparing these findings with those of GWAS and LD analyses, we observed that only two genes, namely, gene_32532 and gene_51857, were shared. Remarkably, these genes both encode pentatricopeptide repeat-containing proteins and were found to be associated with a SNP (position 156 on scaffold5479) identified by both methodologies.

With respect to GO terms, we identified 632 terms associated with the analyzed genes (Supplementary Table S8). The predominant GO term for the cellular component category was ‘nucleus’, which was associated with 168 genes. For the molecular function category, ‘ATP binding’ was the most prominent term and was linked to 89 genes. In terms of biological processes, ‘protein transport’ was associated with 30 genes. The second most prevalent biological process was ‘regulation of transcription, DNA-templated’, which was associated with 27 genes. This finding, in conjunction with the prevalence of ATP binding functions, aligns well with the findings from GWAS and LD analyses.

Furthermore, our analysis revealed insights into carbohydrate-related biological processes. We observed associations with carbohydrate homeostasis (3 genes), the carbohydrate metabolic process (2 genes), and carbohydrate transport (2 genes). This underscores the potential of our approach to identify genes involved in the broader mechanisms of sugar production and storage in sugarcane.

By conducting an enrichment analysis of these genes, we identified 34 GO terms enriched for molecular functions and 39 terms for biological processes (Supplementary Table S10). Among the enriched biological processes, the negative regulation of the transforming growth factor beta receptor signaling pathway, glutathione catabolic process, endoplasmic reticulum membrane fusion, and regulation of phosphate transport were the most significantly enriched processes.

### Differential Expression Analyses

To identify DEGs between IN84-58 (the *S. spontaneum*-representative genotype) and the hybrids SP80-3280 and R570, we developed a gene expression model incorporating development time and genotype as factors. We then compared gene expression levels across genotypes. Our analysis revealed a total of 19,511 DEGs (8,630 upregulated in IN84-58 and 10,881 upregulated in SP80-3280) and 20,869 DEGs (9,338 upregulated in IN84-58 and 11,531 upregulated in R570) when comparing IN84-58 with SP80-3280 (Supplementary Table S12) and R570 (Supplementary Table S13), respectively. Although the differences were not pronounced, the majority of DEGs were downregulated in IN84-58.

To potentially identify DEGs associated with variations in sugar accumulation profiles, we conducted a comparative analysis of the developmental times of the SP80-3280 and R570 genotypes. Specifically, we examined gene expression patterns between 6 and 8 months, 8 and 10 months, and 10 and 12 months for both the SP80-3280 (Supplementary Table S14) and R570 (Supplementary Table S15) genotypes. This comparison aimed to elucidate alterations in sugarcane development possibly linked to processes involved in the interplay between growth and sugar accumulation processes. Our observations revealed distinct profiles between the two genotypes. SP80-3280 exhibited more pronounced differences toward the later stages of development (10 to 12 months), and R570 displayed greater disparities during the earlier stages (6 to 8 months) (Table 4).

**Table 4.** Differentially expressed genes (DEGs) identified through comparisons of development times between the SP80-3280 and R570 genotypes.

| Condition 1 | Condition 2 | Number of DEGs | Upregulated in Condition 1 | Downregulated in Condition 1 |
| --- | --- | --- | --- | --- |
| SP80-3280<br>(6 months old) | SP80-3280<br>(8 months old) | 985 | 593 | 392 |
| SP80-3280<br>(8 months old) | SP80-3280<br>(10 months old) | 465 | 89 | 376 |
| SP80-3280<br>(10 months old) | SP80-3280<br>(12 months old) | 4,746 | 1,157 | 3,589 |
| R570<br>(6 months old) | R570<br>(8 months old) | 2,250 | 569 | 1,681 |
| R570<br>(8 months old) | R570<br>(10 months old) | 2,070 | 122 | 1,948 |
| R570<br>(10 months old) | R570<br>(12 months old) | 1,174 | 753 | 421 |

The total numbers of DEGs identified in the developmental time comparisons between sugarcane varieties SP80-3280 and R570 were 5,559 and 4,936, respectively. Subsequently, we intersected these sets with the DEGs identified from the expression differences between IN84-58 and both SP80-3280 and R570. This analysis revealed 2,649 DEGs for SP80-3280 and 2,468 for R570. To further refine the DEG candidates for investigation alongside the genotype-phenotype associations, we intersected these two sets, resulting in a final set of 853 DEGs. Employing this strategy allowed us to pinpoint a group of DEGs exhibiting differences at various development times between SP80-3280 and R570 and between these two genotypes and IN84-58 (Fig. 2b, Supplementary Table S16). Visualization of the expression patterns of these genes via a heatmap illustrates their contrasting profiles (Fig. 2c).

Our investigation revealed associations between genes identified as DEGs and findings from the other approaches employed (GWAS, LD and FS). Specifically, gene_34382, annotated as an anion transporter, was linked to a SNP identified through a GWAS for Brix and POL traits in Pop1, located at position 210 on scaffold16204. Additionally, gene_71279 and gene_86546, both associated with the transcription factors MYB36, MYB87, RAX1, RAX2, and RAX3, were correlated with LD associations according to GWAS results at position 53 on scaffold196356. Furthermore, gene_10640 (encoding Flavanone 3-dioxygenase 2, Gibberellin 3-beta-dioxygenase 1, and Jasmonate-induced oxygenase), gene_33300 (encoding Flavanone 3-dioxygenase 2 and Jasmonate-induced oxygenase), and gene_52053 (encoding Flavanone 3-dioxygenase 2, Jasmonate-induced oxygenase, and Leucoanthocyanidin dioxygenase) were associated with the SNP at position 50 on scaffold108823 according to the FS approach.

In addition, although not directly linked to the same set of genes, we identified shared annotations between the DEGs and the genotype-phenotype associations. Notably, the FAR1 protein exhibited associations with gene_8664, a DEG identified in our study, and with gene_11104, gene_4861, gene_5373, gene_6529, and gene_8883, all of which were linked to a SNP associated with Brix and POL traits (located at position 111 on scaffold32047) according to the GWAS. Similarly, the E3 ubiquitin-protein ligase showed associations with several DEGs, including gene_96766, gene_99723, gene_112869, gene_25927, gene_31683, gene_76160, and gene_99050, all of which were associated with a SNP detected within the LD set. Furthermore, beta-glucosidase was associated with various DEGs and with gene_50226, which is a gene linked to an LD result (a SNP at position 300 on scaffold413444).

Furthermore, we identified 48 enriched biological process GO terms (Supplementary Table S17). These terms encompass various biological functions, such as defense response (e.g., ethylene-activated signaling pathway, defense response to fungus, response to heat, and response to jasmonic acid), plant development (e.g., gibberellin biosynthetic process and cell wall macromolecule catabolic process), and regulatory processes (e.g., regulation of DNA-templated transcription and induction of programmed cell death). Moreover, we identified 36 distinct enriched molecular function GO terms. Notably, these terms included UDP-glucose 4-epimerase activity and 9-cis-epoxycarotenoid dioxygenase activity. These molecular functions play pivotal roles in processes associated with sucrose accumulation and plant metabolism.

### Gene Coexpression Networks

To comprehensively integrate our findings, we constructed a gene coexpression network employing RNA-Seq gene expression estimates and the WGCNA methodology. Utilizing Pearson correlation coefficients, we computed a gene expression correlation matrix, subsequently fitting the network into a scale-free topology with a β power of 6, yielding an R² value of ∼0.808 and a mean connectivity of ∼992.082. By employing hierarchical clustering, we delineated 250 distinct modules within the network (Supplementary Table S18), ranging from a minimum of 50 genes in group 249 to a maximum of 1,345 genes in group 0. The average gene count per module was approximately 184.40, with a median of 128.5 and a standard deviation of approximately 177.33.

In our investigation, each network group was analyzed for the presence of genes associated with GWAS/LD, FS, or DEGs (Supplementary Table S19). Our findings revealed that 64 groups harbored at least one gene associated with GWAS/LD, 155 groups harbored at least one gene associated with FS, and 146 groups harbored at least one DEG. Notably, 32 groups were concurrently associated with all three approaches. By focusing on these 32 groups and computing the median number of genes per group associated with GWAS/LD, FS, and DEGs, we identified 1, 3, and 8 genes, respectively. We identified and focused our subsequent analysis on groups meeting or surpassing these thresholds, leading to the selection of 8 groups (labeled 0, 2, 9, 12, 15, 18, 36, and 63) for in-depth investigation (4,939 genes).

We performed a GO enrichment evaluation of each of these groups (Supplementary Table S20). Only group 0 presented one biological process term (photosynthesis, light harvesting in photosystem I) enriched according to the established criteria. In relation to molecular function GO terms, group 0 presented three enriched terms (metal ion binding, DNA-binding transcription factor activity, and chlorophyll binding), and group 12 presented two enriched terms (naringenin 3-dioxygenase activity and ATP binding).

Using less stringent criteria (nonadjusted p value of 0.01), we identified additional significant terms associated with sucrose metabolism and related processes. In group 0, the sucrose biosynthetic process (p = 0.00765) and sucrose-phosphate synthase activity (p = 0.00961) were enriched. In group 12, terms related to the response to sucrose (p = 0.00072) and sucrose transport (p = 0.0045) were significantly enriched. Similarly, in group 15, terms related to carbohydrate transport (p = 0.00436), carbohydrate binding (p = 0.00174), and sucrose alpha-glucosidase activity (p = 0.0094) were significantly enriched. In group 18, sucrose transport (p = 0.00165) and sucrose alpha-glucosidase activity (p = 0.00607) were enriched. Additionally, in group 36, sucrose 1F-fructosyltransferase activity (p = 0.00041) was enriched, and in group 63, carbohydrate metabolic processes (p = 0.00258) were enriched.

These findings suggest that, in comparison to other network modules, individual groups within the identified clusters do not exhibit distinct or pronounced specific roles. This lack of specificity arises from the broad impact of their functions across plant metabolism, as many processes performed by these groups are also integral to other modules. However, when all genes within these groups were aggregated and a comprehensive enrichment analysis was conducted (Supplementary Table S21), the enrichment of more biological processes emerged. These enriched processes included the regulation of DNA-templated transcription and positive regulation of the salicylic acid-mediated signaling pathway. These findings imply that the collective action of genes within these groups may exert influence over a range of processes executed by the selected network clusters.

Finally, by leveraging the genes identified within these 8 groups and employing the HRR approach, we constructed three distinct gene coexpression networks: (i) a network tailored to the expression data of the hybrid R570 (Fig. 4a); (ii) a network for the hybrid SP80-3280 (Fig. 4b); and (iii) a network specific to the IN84-58 genotype (Fig. 4c). Considering a total of 4,939 genes, network (i) comprised 2,051 genes and 5,078 edges (with 55 genes having more than 25 connections), network (ii) comprised 2,370 genes and 5,467 edges (with 53 genes having more than 25 connections), and network (iii) comprised 2,791 genes and 7,963 edges (with 112 genes having more than 25 connections). The reduction in gene count is attributed to the HRR methodology, which selectively retains the most robust associations.

**Fig. 4.**
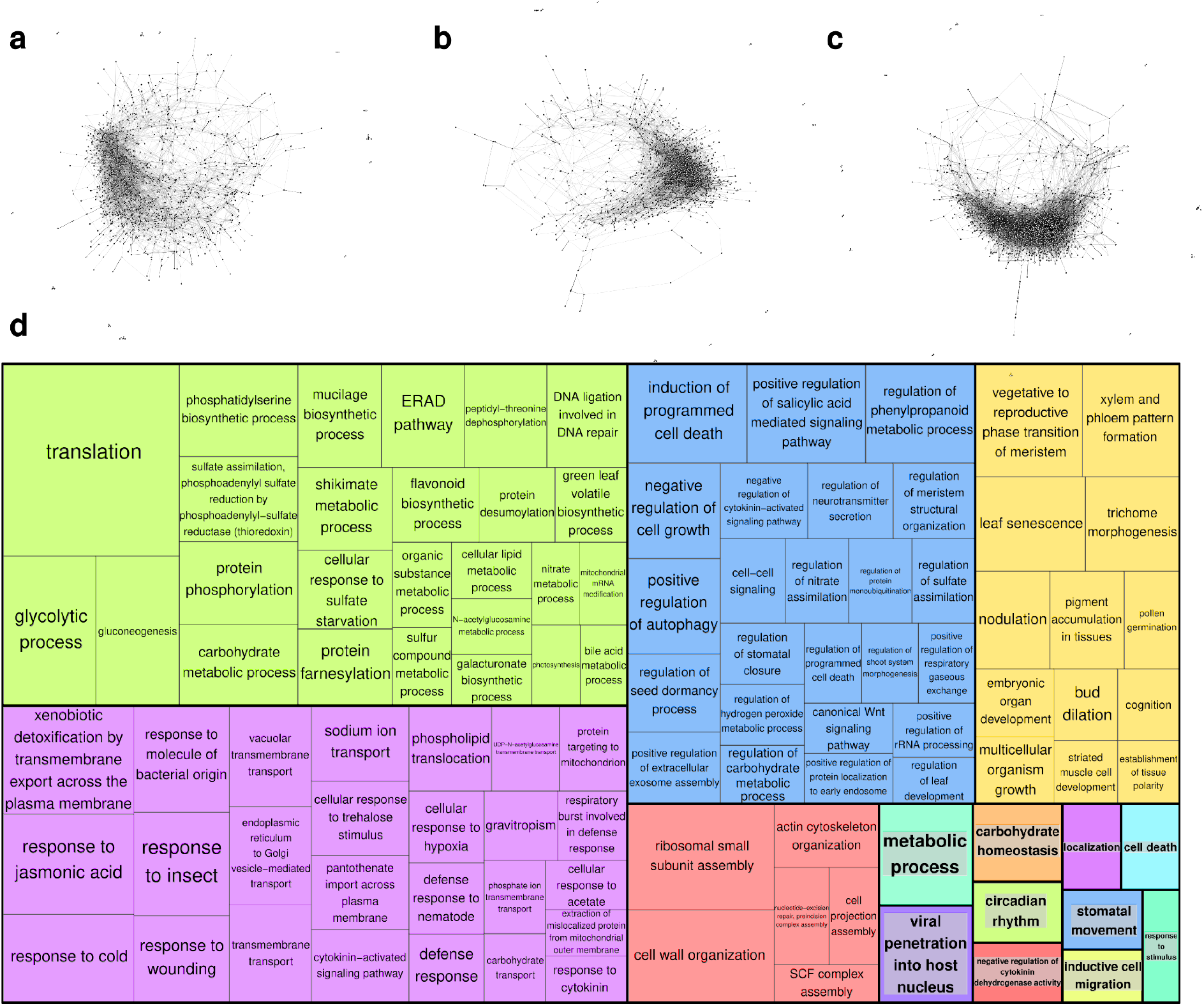
Specific gene coexpression networks modeled using subgroups selected from the original network constructed with the entire set of genes, separated according to (a) sugarcane hybrid R70, (b) hybrid SP80-3280, and (c) IN84-58, a representative genotype of *S. spontaneum*. (d) Gene Ontology (GO) categories associated with biological processes within these selected groups.

This disparity underscores the distinct structural characteristics of the networks, with network (iii) exhibiting a more condensed architecture than networks (i) and (ii). This discrepancy potentially signifies the distinct manners in which the biological functions correlated with these genes are coordinated in each genotype (Fig. 4d), including carbohydrate metabolic processes, carbohydrate transport, and regulation of carbohydrate metabolism. These processes hold significant relevance in the investigation of sucrose accumulation in sugarcane.

Furthermore, given the broad spectrum of biological processes associated with these genes (Fig. 4d) and their potential relevance to sugar accumulation in sugarcane, we examined the gene interactions within each network using centrality measures to pinpoint key genes orchestrating these mechanisms. For each network, we assessed centrality measures, including degree, hub score, and betweenness. A comparison of the network for R570 (Supplementary Table S22), the network for SP80-3280 (Supplementary Table S23), and the network for IN84-58 (Supplementary Table S24) revealed notable disparities in the distribution of gene connections and the identification of pivotal genes driving network structure (Table 5). This observation underscores the distinct regulatory pathways that may lead to the activation of common biological processes in different genotypes.

**Table 5.** Centrality evaluations for specific gene coexpression networks modeled using the highest reciprocal rank (HRR) approach and the genotypes R570, SP80-3280, and IN84-58.

| Network | Statistic | Degree | Kleinberg's Hub Score | Betweenness |
| --- | --- | --- | --- | --- |
| R570 | Minimum | 1 | 0 | 0 |
|  | Maximum | 61 | 1 | ~176011.76 |
|  | Mean | 4.061 | 0.0255393 | 5349.1 |
|  | Median | 2 | 0.0043837 | 395.2 |
|  | Standard Deviation | 6.063621 | 0.06725531 | 14,058.07 |
|  | Top 3 | Nonannotated gene (value of 61); bifunctional aspartokinase/homoserine dehydrogenase 1 (value of 50); and zinc finger FYVE domain-containing protein 26 (value of 50) | Bifunctional aspartokinase/homoserine dehydrogenase 1 (value of 1); GEL complex subunit OPTI (value of ~0.97); and nonannotated gene (value of ~0.95) | Nonannotated gene (value of ~176,011.76); nonannotated gene (value of ~161,517.31); and protein translation factor SUI1 (value of ~145,158.23) |
| SP80-3280 | Minimum | 1 | 0 | 0 |
|  | Maximum | 62 | 1 | ~184,912.91 |
|  | Mean | 4.614 | 0.062281 | 4,841.1 |
|  | Median | 2 | 0.022610 | 526.8 |
|  | Standard Deviation | 6.294851 | 0.09964838 | 12,779.9 |
|  | Top 3 | Large ribosomal subunit protein eL18 (value of 62); transcription factor BTF3/basic transcription factor 3 (value of 55); and Calcium/calmodulin-regulated receptor-like kinase 1 (value of 46) | Senescence associated gene 20 (value of 1); Large ribosomal subunit protein eL18 (value of ~0.99); and nonannotated gene (value of ~0.94) | Transcription factor BTF3/basic transcription factor 3 (value of ~184,912.91); large ribosomal subunit protein eL18 (value of ~166,457.51); and Calcium/calmodulin-regulated receptor-like kinase 1 (value of ~113,026.88) |
| IN84-58 | Minimum | 1 | 0 | 0 |
|  | Maximum | 51 | 1 | ~171,446.33 |
|  | Mean | 5.706 | 0.060052 | 5033.2 |
|  | Median | 3 | 0.024053 | 644.1 |
|  | Standard Deviation | 7.503239 | 0.09622554 | 11,460.91 |
|  | Top 3 | Adagio protein (value of 51); glutathione S-transferase (value of 49); and nonannotated gene (value of 48) | Acyl-coenzyme A thioesterase 13 (value of 1); cinnamoyl-CoA reductase 1 (value of ~0.97); and auxin response factor (value of ~0.90) | Kinetochore-associated protein KNL-2 (value of ~171,446.33); formin-like protein (value of ~90,953.78); and protein Weak Chloroplast Movement Under Blue Light (value of ~85,236.03) |

## Discussion

### Strategies for Dealing with Sugarcane Genetic Complexity

In our study, we employed various innovative strategies to overcome the genomic intricacies of sugarcane in order to investigate the molecular basis of the most relevant trait of this crop. One of the primary obstacles encountered when investigating sugarcane polymorphisms are aneuploidies, which manifest as variable numbers of alleles per chromosome and distinct genomic regions harboring different allele copy numbers within the same chromosome (Zhang et al. 2018; Aono et al. 2021).

Traditionally, addressing such complexity has involved either simplifying SNP markers by assuming fixed ploidy (Fickett et al. 2019; Yang et al. 2020; Pimenta et al. 2021; Wang et al. 2023a; Zhang et al. 2023) or estimating specific ploidy levels for individual markers (Balsalobre et al. 2017; Batista et al. 2022). However, in our study, rather than disregarding allele variations, we opted to represent SNPs not as dosages but as allele proportions. This approach allowed us to retain a significant number of markers that would otherwise have been discarded due to the low statistical power of the dosage estimation process (Aono et al. 2020).

In recent years, there have been notable advancements in sugarcane genomics, with the emergence of several genomic references, including those tailored for allele specificity (Zhang et al. 2018; Bao et al. 2024; Healey et al. 2024). Although these resources have significantly enhanced sugarcane genomic studies, accurately aligning short sequencing reads to these references and inferring correct allele dosages remains a challenge. The sugarcane genome is characterized by a high degree of duplication, leading to a substantial proportion of reads being mapped in duplicate across its genome. The conventional approach to address this issue involves excluding duplicate mapped reads, which, unfortunately, results in a significant reduction in the number of generated SNPs (Gardiner et al. 2016).

However, Aono et al. (2020) demonstrated that this reduction can be circumvented by utilizing a sugarcane methyl-filtered reference (Grativol et al. 2014), which is compatible with the GBS approach employed. Thus, we chose to utilize this reference for SNP calling, thereby overcoming the reduction in SNP numbers observed with other genomic references. Furthermore, to indirectly associate our findings with the genomic references of *S. officinarum* and *S. spontaneum*, we conducted comparative alignments between RNA-Seq-based assembled genes and the methyl-filtered genome scaffolds. By employing this strategy, we not only increased the number of markers but also enhanced the likelihood of identifying associations with QTL regions.

Our approach to addressing the complexity of sugarcane genetics diverged from traditional QTL mapping methods based on linkage analyses. Instead, we employed marker‒trait association tests in Pop2. The current methodologies available for handling polyploid species via linkage analysis do not adequately address the nuances of sugarcane genetics (Mollinari et al. 2020). The construction of linkage maps in sugarcane typically yields numerous unsaturated linkage groups characterized by substantial intermarker distances (Costa et al. 2016; Balsalobre et al. 2017; Yang et al. 2018; You et al. 2019; Wang et al. 2022b; Wang et al. 2022b, 2023a). As a consequence, many markers are excluded from the analysis, thereby limiting the pool of SNPs available for QTL identification. Moreover, we leveraged machine learning approaches to enhance the reliability of our findings. Through the integration of data from two distinct populations and the utilization of diverse methodological strategies, we strengthened the robustness of our inferences.

The exploration of genotype‒phenotype relationships in sugarcane, in conjunction with other omics approaches, is still in its early stages. Only a limited number of studies have investigated these associations within a multiomic framework (Li et al. 2023; Pimenta et al. 2023). We believe that integrating such methodologies has greatly enhanced our ability to infer the underlying biological mechanisms governing sucrose accumulation in sugarcane. Although the fundamental mechanisms of sucrose metabolism are widely acknowledged (Sachdeva et al. 2011; Datir and Joshi 2016), the factors contributing to enhanced sucrose accumulation remain incompletely understood. Consequently, integrating the findings from various omics analyses, particularly through coexpression analysis, has provided a comprehensive and valuable dataset.

### Novel Insights into Sugarcane Sucrose Accumulation

Sugarcane is the crop with the greatest capacity for sucrose storage (Qin et al. 2021). Consequently, breeding programs for sugarcane have prioritized the development of varieties with optimized sucrose storage capabilities. Variations in sucrose content within sugarcane varieties are attributed to a complex interplay of polygenic effects, diverse biological processes and environmental effects (Khan et al. 2023). Previous GWASs have elucidated the association of sucrose accumulation with polymorphisms located near different genes. These genes encompass annotations mostly related to plant growth, development (Racedo et al. 2016; Fickett et al. 2019; Wang et al. 2023b), and responses to both biotic and abiotic stresses (Wang et al. 2023b; Zhang et al. 2023). In our GWAS analysis, although we found noteworthy similarities with previous studies, particularly regarding the involvement of phosphatases, kinases, and ubiquitin-like proteins (Fickett et al. 2019; Wang et al. 2023b; Zhang et al. 2023), we were able to expand upon these findings.

The involvement of sucrose signaling pathways in regulating various growth and developmental processes is widely recognized in the literature (Papini-Terzi et al. 2009; Chen et al. 2019). Moreover, the intricate interplay between sucrose and plant hormones, such as abscisic acid, salicylic acid, jasmonic acid, and ethylene, underscores the multifaceted nature of the association between sucrose and stress responses. Sucrose serves as an energy source to cope with stress, and at different levels, it plays pivotal roles in regulating the expression of stress-responsive genes (Khan et al. 2023).

Our investigation, supported by the literature, underscores the synergistic mechanism wherein sucrose levels impact stress response and growth dynamics. Notably, for Pop1, we identified GWAS-associated SNPs surrounding genes annotated for anion transporters, FAR1 proteins, and serine/threonine-protein kinases. These genes play pivotal roles in balancing growth and stress responses (Zheng et al. 2010; Ramesh et al. 2015; Liu et al. 2019; Jiang et al. 2022) and have potential implications for carbohydrate synthesis (Ma et al. 2017; Luo et al. 2020; Liu et al. 2022). Furthermore, our GWAS of Pop2 revealed a gene annotated for a pentatricopeptide repeat-containing (PPR) protein, which has also been implicated in both plant development and stress response pathways (Liu et al. 2017; Pimenta et al. 2023). Moreover, PPR proteins are implicated in the modulation of gene expression in organelles and play crucial roles in plant embryogenesis (Cushing et al. 2005; Yin et al. 2013), potentially accounting for the observed enrichment of GO terms associated with embryonic development.

Although the use of the sugarcane methyl-filtered genome reference enabled us to detect a significantly greater number of SNPs, the assessment of LD decay patterns was hindered by the fragmented nature of this assembly. Nevertheless, broadening the analysis to include LD associations with GWAS-identified markers across the entire SNP set, irrespective of their scaffold location, allowed us to retrieve a more extensive set of genes, thereby facilitating more comprehensive inferences.

Consistent with our GWAS findings, we also identified additional genes associated with stress responses in the LD associations. These include E3 ubiquitin-protein ligase (Shu and Yang 2017), calcineurin B-like protein 10 (Su et al. 2020), RING finger protein 141 (Han et al. 2022), abscisic acid 8’-hydroxylase 2 (Umezawa et al. 2006), DEAD-box ATP-dependent RNA helicase 25 (Kim et al. 2008), and peroxisomal biogenesis factor 3 (Hu et al. 2012). Notably, several stress-responsive genes are associated with sucrose accumulation, potentially leading to changes in carbon allocation and photosynthetic activities (Verma et al. 2019; Qin et al. 2021).

Additionally, through LD expansion, we successfully identified key players involved in sucrose synthesis and accumulation. Our analysis revealed genes associated with crucial processes, including bZIP transcription factor, beta-glucosidase, and thioredoxin-like protein genes. The bZIP transcription factor has previously been recognized as a negative regulator of cold and drought responses in rice (Liu et al. 2012). It also plays a significant role in various carbohydrate-associated processes, highlighting the intricate relationship between stress responses and growth dynamics. Moreover, in addition to its involvement in starch regulation in rice (Wang et al. 2013a), bZIP has been implicated in sucrose synthesis, transport, and metabolism (Ma et al. 2019; Stein and Granot 2019), and its role has already been investigated in sugarcane (Wang et al. 2022a).

Furthermore, the beta-glucosidase protein has been linked to sucrose synthesis and accumulation (Khan et al. 2023), potentially exerting a negative influence on sucrose accumulation (Qin et al. 2021). Last, thioredoxin (TRX) proteins are associated with trehalose synthesis (Khan et al. 2023), which has been shown to impact sucrose metabolism (De Oliveira et al. 2022). TRX proteins play a pivotal role in modulating chloroplast functions to maintain equilibrium in photosynthetic reactions through redox regulation (Nikkanen and Rintamäki 2019). Consequently, these proteins are intricately linked to carbohydrate metabolism and responses to oxidative stress. Moreover, TRX has previously been identified as a regulator of carbon-nitrogen partitioning in tobacco (Ancín et al. 2021). Overexpression of TRX leads to the accumulation of nitrogen-related metabolites while decreasing carbon-related metabolites.

Even with the LD approach employed alongside GWAS results, we did not identify a significant number of genes directly regulating sucrose metabolism, such as sucrose-synthesizing and hydrolyzing enzymes (Datir and Joshi 2016). The lack of further associations related to sucrose metabolism, including sucrose synthase, sucrose phosphate synthase, and invertases, may be attributed to various factors. First, the genes identified through GWAS and LD analyses might exert an indirect influence on these processes, triggering mechanisms that ultimately impact the efficiency of sucrose accumulation through pathways yet to be elucidated, thus warranting further investigation. This is particularly noteworthy in light of previous unsuccessful endeavors to manipulate genes directly linked to sucrose transport and metabolism (Qin et al. 2021).

Moreover, the reduced number of individuals employed in Pop1 for GWAS might have influenced our findings. Although the sucrose content profiles of the selected individuals exhibited high variability, as evidenced by the high heritability estimates of 0.89 and 0.9 for Brix and POL, respectively, increasing the number of genotypes could enhance the observed results. This expansion could facilitate the identification of additional associations, potentially capturing effects with reduced impact on phenotypic variance and lower allele frequencies (Korte and Farlow 2013).

Additionally, the use of GBS has limited our ability to sample various genomic regions for evaluation. Although GBS has the potential to identify a significant number of markers associated with QTLs (Elshire et al. 2011), its coverage of the entire genome is incomplete. Coupled with our employment of a fragmented genomic reference, several regions of the sugarcane genome remained unassessed. Therefore, the utilization of scalable and high-quality long-read sequencing holds great promise for advancing sugarcane genomics, particularly for enabling proper application of the current allele-specific genomic references (Zhang et al. 2018; Bao et al. 2024; Healey et al. 2024).

When evaluating the enriched GO terms associated with the GWAS and LD results, it was possible to observe molecular functions and biological processes primarily pertaining to regulatory activities, such as kinase activity, intracellular transport, and functions related to RNA and DNA processing. Specifically, certain terms are associated with sugar metabolism and the hormone abscisic acid (ABA), which plays a pivotal role in plant metabolism, particularly in response to abiotic stress. Previous investigations conducted on sugarcane have indicated a potential overlap between sugar and ABA-related processes. This overlap arises from the capacity of ABA to regulate a set of genes associated with sucrose metabolism (Papini-Terzi et al. 2009).

In addition to the findings obtained from GWAS, we employed machine learning approaches, a strategy that has proven effective in uncovering genotype‒phenotype associations (Aono et al. 2020; Pimenta et al. 2021, 2023). Through this integrative approach, we present a comprehensive analysis that extends beyond conventional GWAS findings. This enables us to uncover a wider set of metabolic pathways that may be associated with genes implicated in sucrose accumulation.

Our analysis revealed an expanded repertoire of enriched GO terms in the FS results, reflecting a diverse range of regulatory and nonspecific processes. These include posttranslational modifications in proteins, DNA and organelle processing, embryonic development, transport, and nutrient responses. Notably, processes related to growth, hormone signaling, stress responses, and lipid metabolism were also indicated. To date, there has been no direct association between these processes and sugar metabolism documented in the literature. However, it is plausible that, similar to the mechanism associated with ABA, these processes may exert an indirect influence on this process.

When comparing different genotypes, the observed DEGs were implicated in a broad array of biological processes. Thus, when comparing the IN84-58 *S. spontaneum* genotype with the SP80-3280 and R570 hybrid genotypes, subset selection was necessary to identify potential associations with sucrose accumulation profiles. Although sucrose synthesis primarily occurs in sugarcane leaves, sucrose is transported through the phloem to culms, where it is utilized for plant growth and development or is stored (Mason et al. 2020). When the plant reaches maturation, sugars are directed toward storage, accompanied by the activation of specific mechanisms, resulting in changes in accumulation efficiency within the culms (Wang et al. 2013b).

Thus, we selected DEGs between *S. spontaneum* and the hybrids only if they were also detected during contrasting developmental stages. This decision stems from the fact that the gene expression patterns in sugarcane tissues are significantly influenced by the developmental stage (Wang et al. 2013b; Chen et al. 2019). In addition to developmental differences, there are also genotype-specific DEGs (Papini-Terzi et al. 2009). As our focus did not include the specific mechanisms of R570 and SP80-3280, we opted for an intersection between the results obtained from both comparisons, thereby enhancing the reliability of associating such expression changes with sucrose accumulation.

The intersection of the DEG sets led to the identification of 853 genes, revealing intriguing insights. Notably, these genes are associated with biological processes that overlap with those identified through GWAS and FS-selected markers. Regulatory mechanisms involving protein modifications, transcription factors, responses to oxidative stress, anion transport, and DNA/RNA processing were indicated. Additionally, these genes play roles in the response to both biotic and abiotic stresses, with implications for ethylene and gibberellin regulation. Furthermore, associations with sugar catabolism were discerned. This convergence of mechanisms across multiple omics layers underscores the interconnectedness of biological processes and the potential for integrated analyses to increase our comprehension of complex traits.

As anticipated, our analysis revealed genes that exhibited both differential expression and associations with phenotype‒genotype relationships. Among these genes, the only gene that overlapped with GWAS findings was annotated as an anion transporter, reinforcing the potential involvement of its activity in sucrose accumulation. With respect to genes associated with FS-selected markers, we identified one gene encoding the transcription factor MYB36, which has been previously implicated in plant growth and stress response (Monje-Rueda et al. 2023). Additionally, we detected a gene annotated for jasmonate-induced oxygenase, known for its role in suppressing plant immunity (Caarls et al. 2017), providing further insights into the molecular mechanisms underlying disease susceptibility in high Brix genotypes.

Additionally, we also found common annotations between the set of DEGs and the GWAS results. Although they do not correspond to the same genes, it is clear that the same biological mechanisms are associated with phenotypic variability favoring sucrose accumulation and differential gene expression in different sugar content genotypes. The regulatory roles of the FAR1 protein, E3 ubiquitin-protein ligase, and beta-glucosidase warrant further attention because they are implicated in carbohydrate synthesis and potentially influence the balance between sucrose accumulation and the defense response (Ma et al. 2017; Shu and Yang 2017; Liu et al. 2019; Qin et al. 2021; Khan et al. 2023).

While only a limited number of genes were consistently identified across all approaches and datasets, there is a clear consensus emerging regarding the biological processes and mechanisms influenced by these selected genes. To consolidate our findings, we constructed a gene coexpression network. Specifically, our analysis enabled us to delineate eight distinct gene groups within the network comprising DEGs as well as genes exhibiting significant associations with SNPs linked to divergent sucrose accumulation levels, as identified through GWAS and FS.

Based on the premise that the selected genes are correlated with sucrose accumulation, we hypothesize that the most significant differences in the impact of these genes on sucrose accumulation are attributable to their interactions. Therefore, investigating these interactions might provide valuable insights into key genes that could serve as focal points for more extensive investigations. Thus, we constructed specific gene coexpression networks, differentiating between the gene expression profiles of hybrids and the *S. spontaneum* genotype.

The network constructed for *S. spontaneum* gene expression exhibited approximately 50% more connections than the hybrid genotype networks. This suggests that a greater number of gene interactions are necessary for *S. spontaneum* to perform the same biological processes as the hybrids. We believe that the simpler network structure observed in the hybrids signifies more efficient regulation of the processes related to sucrose accumulation through gene interactions. However, external factors, such as stressors, can easily influence gene interactions in the hybrid networks. In contrast, gene communication in *S. spontaneum* is less susceptible to disruption, consistent with the inherent resistance of this species to different types of biotic and abiotic stresses.

While conducting a comprehensive analysis of all network components could provide valuable insights into sucrose accumulation, our study prioritized key network elements. We achieved this by evaluating specific centrality measures, aiming to correlate node influence with the biological implications of gene roles, thus enabling meaningful inferences (Wang et al. 2022c). Furthermore, by comparing genes with high centrality measures across the networks modeled, we can infer differences in the regulatory mechanisms governed by the gene sets within these networks.

Starting with evaluations of degree, which measures the importance of a gene based on the number of connections it possesses, and Kleinberg’s hub score, which incorporates gene proximity to other network nodes into the assessment, it becomes evident that genes exhibiting increased centralities in the hybrid networks are more closely associated with the regulation of fundamental cellular processes crucial for plant growth, including amino acid biosynthesis, signal transduction, gene expression regulation, and protein synthesis. Conversely, in the *S. spontaneum* network, these genes appear to be involved in a broader array of mechanisms, potentially including roles in stress-response signaling pathways, as indicated by glutathione S-transferase (Vaish et al. 2020), adagio protein (Bulgakov et al. 2017), auxin response factor (Li et al. 2016), acyl-coenzyme A thioesterase (Kalinger et al. 2020), and cinnamoyl-CoA reductase 1 (Park et al. 2017). These findings support our observation regarding the association of this network architecture with the effective response of *S. spontaneum* to various types of stress.

Betweenness centrality exhibited an opposite pattern. In the networks modeled for the hybrids, genes with high betweenness were mostly associated with protein synthesis and gene expression regulation, including the protein translation factor SUI1 (Li et al. 2022), the transcription factor BTF3 (Pruthvi et al. 2017), and the calcium/calmodulin-regulated receptor-like kinase 1 (Yuan et al. 2022). In contrast, the network modeled for *S. spontaneum* had genes with high betweenness primarily associated with cellular structure and division, such as the kinetochore-associated protein KNL-2 (Zuo et al. 2022) and formin-like protein (Kollárová et al. 2021). A high betweenness measure indicates that a gene permeates many gene associations, potentially facilitating the flow of interactions within the network. This suggests that in *S. spontaneum*, gene associations favor the maintenance of cellular architecture integrity. Conversely, in hybrid networks, these genes are more involved in signal transduction.

Remarkably, the observed network dynamics suggest that gene communication within the gene set associated with *S. spontaneum* is predominantly associated with plant immunity. In contrast, in the hybrid networks, we observed indications of a more nuanced interplay, potentially influenced by external factors. These findings highlight the intricate regulatory networks underlying sucrose accumulation, revealing distinct regulatory strategies adopted by different genotypes in response to environmental stimuli.

## Conclusion

Sugar production is the primary focus of sugarcane breeding, and this process is governed by complex interactions among polygenic effects and diverse biological processes. Unraveling the genotype‒phenotype associations that significantly increases sucrose content presents a great challenge but holds immense value for sugarcane breeding. Despite these efforts, the development of varieties optimized for this trait remains limited. Genetic modifications targeting genes specific to sucrose metabolism have not yielded the desired outcomes. Thus, comprehensive investigations spanning a broad set of mechanisms are essential for identifying promising targets.

In our study, we adopted an integrative approach to examine sugarcane genetics. By combining GWAS, machine learning algorithms, and differential expression analyses, we identified key factors involved in sucrose accumulation that warrant attention. Notably, a jasmonate-induced oxygenase was identified as a DEG associated with significant findings from our GWAS. The mutation observed near this gene, known for its role in suppressing plant immunity, appears to favor sugar accumulation. Additionally, the role of the beta-glucosidase protein was noteworthy, with annotations found in genes proximal to GWAS hits and DEGs. Given its negative impact on sucrose accumulation, this enzyme is a promising target for biotechnological investigations.

Moreover, we integrated all genes associated with our findings across analyses and datasets into a comprehensive gene coexpression network, providing a foundation for future genetic studies. Contrasts between specific gene coexpression networks constructed for *S. spontaneum* and sugarcane hybrids revealed differences in gene associations linked to sugar accumulation. We hypothesize that the simpler network structure observed in hybrids may indicate a more efficient process, albeit potentially more susceptible to external influences such as stressors. Conversely, the more cohesive network observed in *S. spontaneum* may be associated with enhanced plant immunity.

## Statements and Declarations

### Author Contributions

**Alexandre Hild Aono**: Conceptualization, Data Curation, Methodology, Software, Validation, Visualization, Formal analysis, Investigation, Writing - Original Draft, Writing – review & editing. **Ricardo José Gonzaga Pimenta**: Conceptualization, Data Curation, Investigation, Writing - Review & Editing. **Jéssica Faversani Diniz**: Investigation, Writing - Original Draft. **Marishani Marin Carrasco**: Investigation. **Guilherme Kenichi Hosaka**: Data curation, Formal analysis, Writing – review & editing. **Fernando Henrique Correr**: Data curation, Formal analysis, Writing – review & editing. **Ana Letycia Basso Garcia**: Formal analysis. **Estela Araujo Costa**: Data curation. **Alisson Esdras Coutinho**: Data curation. **Luciana Rossini Pinto**: Data curation, Funding acquisition, Resources, Supervision. **Marcos Guimarães de Andrade Landell**: Data curation, Funding acquisition, Resources. **Mauro Alexandre Xavier**: Data curation, Funding acquisition, Resources. **Dilermando Perecin**: Data curation, Supervision. **Monalisa Sampaio Carneiro**: Data curation, Funding acquisition, Resources. **Thiago Willian Balsalobre**: Data curation. **Reginaldo Massanobu Kuroshu**: Data curation, Formal analysis, Supervision, Validation, Writing – review & editing. **Gabriel Rodrigues Margarido**: Data Curation, Formal analysis, Funding acquisition, Resources, Supervision, Validation, Writing – review & editing. **Anete Pereira de Souza**: Conceptualization, Funding acquisition, Project administration, Resources, Supervision, Writing – review & editing.

### Data Availability

The accession codes of the sequencing data are available through the Sequence Read Archive (SRA) database with the accession numbers PRJEB40481, PRJNA702641, and SRP151376.

